# Algal Optics

**DOI:** 10.1101/2025.06.18.660391

**Authors:** Ming Yang, Sumit Kumar Birwa, Raymond E. Goldstein

## Abstract

Nearly a decade ago it was discovered that the spherical cell body of the alga *Chlamydomonas reinhardtii* can act as a lens to concentrate incoming light onto the cell’s membrane-bound photoreceptor and thereby affect phototaxis. Since many nearly transparent cells in marine environments have complex, often non-axisymmetric shapes, this observation raises fundamental, yet little-explored questions in biological optics about light refraction by the bodies of microorganisms. There are two distinct contexts for such questions: the *absorption* problem for *incoming* light, typified by photosynthetic activity taking place in the chloroplasts of green algae, and the *emission* problem for *outgoing* light, where the paradigm is bioluminescence emitted from scintillons within dinoflagellates. Here we examine both of these aspects of “algal optics” in the special case where the absorption or emission is localized in structures that are small relative to the overall organism size, taking into account both refraction and reflections at the cell-water boundary. Analytical and numerical results are developed for the distribution of light intensities inside and outside the body, and we establish certain duality relationships that connect the incoming and outgoing problems. For strongly non-spherical shapes we find lensing effects that may have implications for photosynthetic activity and for the angular distribution of light emitted during bioluminescent flashes.

## I. INTRODUCTION

In a remarkably prescient paper [1], Kessler, Nedelcu, Solari and Shelton found in 2015 that the spheroidal cell bodies of various green algae can act as lenses, bringing light from a distant source to a focus outside the cell. They suggested there might be functional significance to such a lensing effect, and indeed in the following year Ueki, *et al*. [2] found the first example, using the green alga *Chlamydomonas reinhardtii*. Phototaxis in *Chlamydomonas* is achieved by the coupling between signals received by a photosensor and the beating dynamics of the two flagella that are anchored below the cell wall near the anterior pole of the cell. The photosensor (Fig. 1) sits within a membrane at the periphery of the cell, and in wild type cells has behind it an “eye spot”, a pigmented protein layer visible in bright field microscopy. Acting as a quarter-wave plate [3], the eye spot reflects incident light and thus blocks light from behind from the photo-sensor, such that only light coming from outside the cell is detected. It is precisely this directionality that underlies the ability of cells to steer toward or away from light [4, 5]. For example, in negatively phototactic mutants lacking the eye spot, light from behind the cell falls on the photosensor, and one might naively expect the cell to be unable to perform phototaxis because of the isotropic detection of light. Yet, because of the lensing effect of the cell the intensity of light falling on the photoreceptor from behind is greater than that from the forward direction, giving rise to a distinguishable signal and thus to net phototactic motion (erroneously) toward the light. It is thus plausible that the eyespot was selected by evolution precisely because, in providing directionality to the sensing of light, it improves the phototaxis of cells [6].

**FIG. 1.**
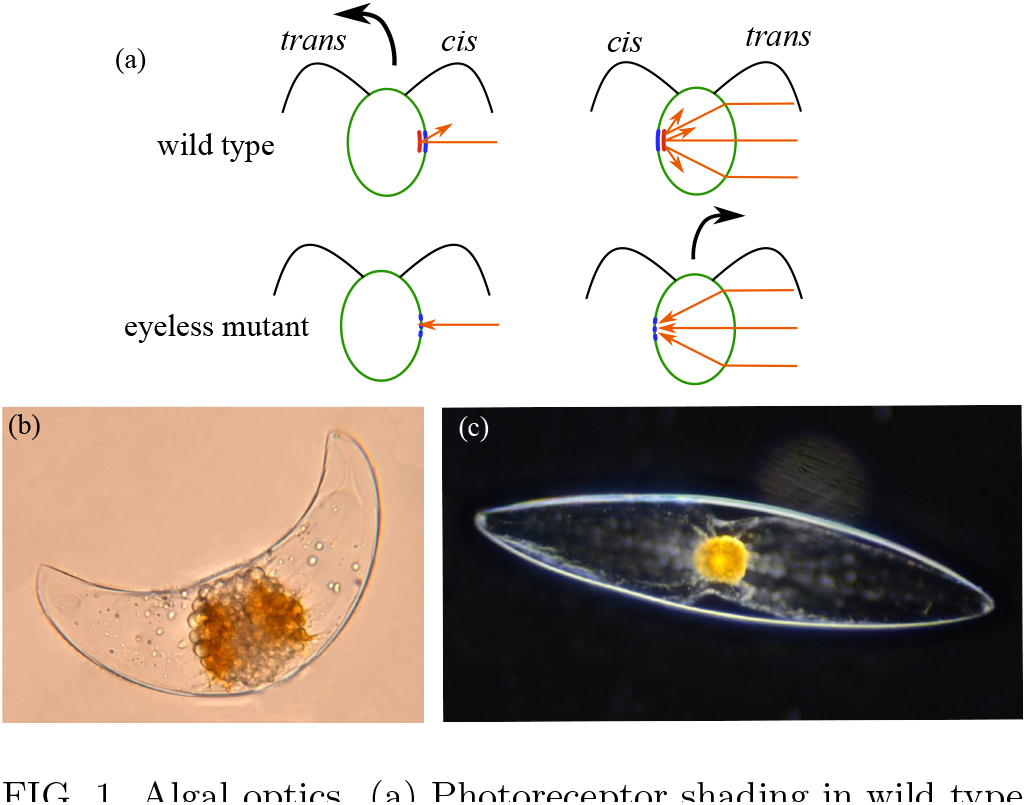
Algal optics. (a) Photoreceptor shading in wild type and eyeless mutants of *Chlamydomonas reinhardtii*, after [2]. Dinoflagellates: (b) crescent-shaped *Pyrocystis lunula* and (c) spindle-shaped *Pyrocystis fusiformis* [9].

Lenses and lens-like properties of cells are known in other systems. For example, there is evidence for a lens *within* the eyespot of the dinoflagellate *Nematodinium* [7] that helps to gather light onto its photoreceptor. At the same time as the study of Ueki, *et al*., separate work on cyanobacteria [8] showed that its cell body can act as a lens, creating an uneven distribution of light intensity in the inner cell wall. Further work showed that a type of supermolecular machine for cell mobility is activated by that directional light [10]. These kinds of effects are now broadly understood for prokaryotes [11]. Finally, we note the case of diatoms, algae whose surface has a complex silica microstructure patterned on the scale of the wavelength of visible light. These structures induce wavelength-dependent optical properties [12, 13] and diffraction [14].

Taking a broader view, there are important historical examples where lensing effects are thought to impact on life’s processes, particularly with regard to plants [15]. For example, phototropism, the movement of plants toward light, is dependent on the fact that the cytoplasm of plant cells has a higher index of refraction than the surrounding medium (air or water). This was demonstrated by an experiment [16] that showed that this movement reverses direction when a plant is immersed in a medium with higher index of refraction than the cytoplasm. Many investigations have focused on the effect of cell shape on photosynthesis in plants and fungi. For example, a study [17] of the filamentous fungus *Phycomyces blakesleeanus* showed that the light intensity at the cell surface is enhanced by a factor of ∼ 2, while theoretical work suggests an even larger boost [18]. The epidermal cells of certain tropical plant species are thought to act as lenses, providing an advantage in light-gathering ability for shade plants [19], although later work found that the structure does not help to gather diffuse light [20].

The results summarized above indicate that lensing effects by the cell bodies of microorganisms can have a functional significance, even in the simplest possible geometry of a sphere. Yet, there are many freshwater and marine microorganisms with strikingly non-spheroidal shapes, as exemplified by dinoflagellates such as *Pyrocystis lunula* and *Pyrocystis fusiformis* shown in Fig. 1(b,c). The properties of such complex, often non-axisymmetric bodies appear to be completely unknown in the biological optics literature, although there is now a growing field of “freeform optics” [21–24] that considers nonaxisymmetric shapes. Motivated by these strange and wondrous forms, in this paper we commence an in-depth study of the field of “algal optics”.

Beyond their phototaxis, organisms such as green algae and dinoflagellates are photosynthetic and their chloroplasts, the organelles containing the photosynthetic apparatus, are in characteristic positions within the cell. In the case of dinoflagellates, the chloroplasts move around the cell in diurnal patterns [25–27], changing the absorption profile of the cytoplasm [28]. A natural question is whether lensing can enhance the intensity of light falling on chloroplasts. This is the “incoming” problem.

Dinoflagellates are among the many marine and fresh-water organisms that exhibit bioluminescence [29]. Unlike the steady glow of bioluminescent bacteria, these eukaryotes emit bright flashes of light in response to fluid or mechanical shear [30, 31]. This light emanates from membrane-enclosed organelles termed “scintillons”, within which occur chemical reactions involving the protein luciferin. A second natural question is thus whether lensing can alter the spatial distribution of light emitted from such sources. This is the “outgoing” problem.

Figure 2 presents ray-tracings using a ray optics simulator [32] that illustrate how light is emitted from internal sources in different dinoflagellate cell shapes. The geometries represent the spindle-shaped cell body of *P. fusiformis* (a,b) and the crescent-shape of *P. lunula* (c,d).

**FIG. 2.**
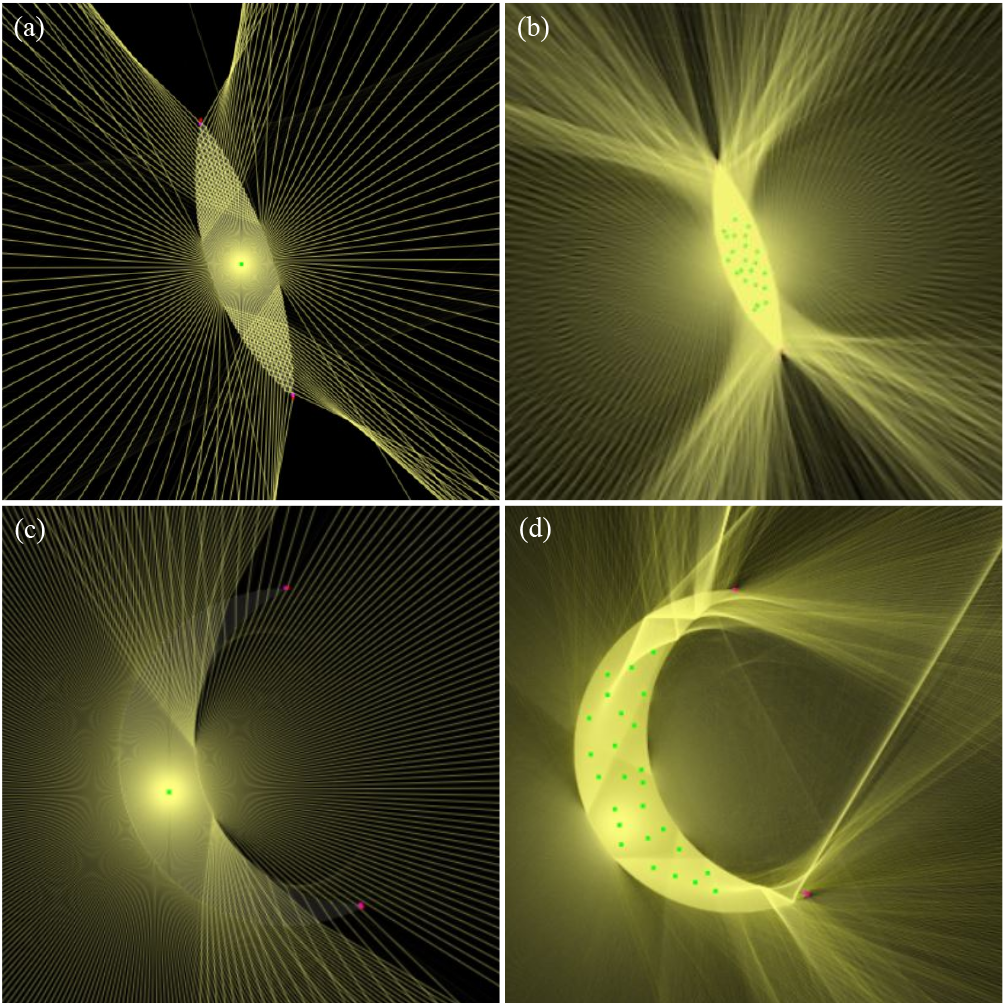
Ray tracings for various dinoflagellate geometries [32]. Single and multiple sources for the shape of *P. fusiformis* (a,b) and of *P. lunula* (c,d).

The green dots indicate the positions of bioluminescent sources within the cell body. In (a,b) the emitted rays are symmetrically distributed about the long axis, with intensity primarily directed laterally—toward the left and right—and little light escaping along the axial (top–bottom) directions. This symmetry and directional focus suggest that the fusiform shape acts as a cylindrical lens, favoring horizontal emission.

In the single-source case for the crescent shape (c), the emission exhibits a markedly different pattern; light is preferentially emitted along the axial directions (top and bottom), forming focused streaks of high intensity. Moreover, there is an enhanced concentration of rays on the concave side of the cell, particularly close to the cell body, due to the curvature-induced refraction. In the multiple-source case (d), the overall emission appears more spatially diffuse, but the asymmetry persists, with noticeably more rays directed toward the concave side. The convex side emits less light overall, likely due to the outward bend redirecting rays away from those regions. These results highlight the critical role of geometry in shaping the angular emission profile. Even in the absence of specialized optics, the cell’s shape can redistribute internally generated light in a highly non-uniform manner.

In this paper we study the incoming and outgoing problems with analytical and numerical methods. For the motivational cases of green algae and dinoflagellates there is a reasonable separation between the size of the absorber or emitter (the algal photoreceptor is ∼ 1 ‒ 2 *µ*m across, scintillons are ∼0.5 − 1.5 *µ*m in diameter) and the size of the entire cell (*Chlamydomonas* is ∼ 10 *µ*m cross, dinoflagellates can be ∼ 150 *µ*m long). Given this, in the simplest model we consider absorbers/emitters to have a radius *a* ≪ *R*, where *R* is a characteristic size of the cell. Section II sets up the incoming and outgoing problems in mathematical terms, laying out the additional modeling assumptions and definitions. As an example of the problem of interest, the observations of Ueki, et al. are analyzed quantitatively to gain insight into the lensing effects that can occur. Analytical results for two-dimensional bodies are presented in § III, including the results of averaging over orientations. Three-dimensional problems are considered analytically in § IV and numerically in §V, where we illustrate strong, complex lensing effects associated with shapes like those of the genus *Pyrocystis*. In §VI we discuss possible experiments to examine this problem in greater detail.

## II. PRELIMINARIES

We consider two complementary problems, each motivated by considerations of the natural environment.

### The incoming problem: How cell geometry affects light absorption in photosynthesis

As shown in Figure 3(a), this problem explores how a curved cell wall modifies the light intensity at a specific location inside the cell. In the turbulent ocean, microorganisms receive sunlight that is scattered and refracted from nearly all directions, while their random orientations within the flow further homogenize the light distribution. We consider the case in which the cell geometry and random flows are such that there is a uniform angular distribution of cell orientations, so the incoming light appears isotropic from the cell’s perspective. The mathematical problem of interest is then the light intensity received by a small target within the cell relative to that in the absence of the surrounding cell.

**FIG. 3.**
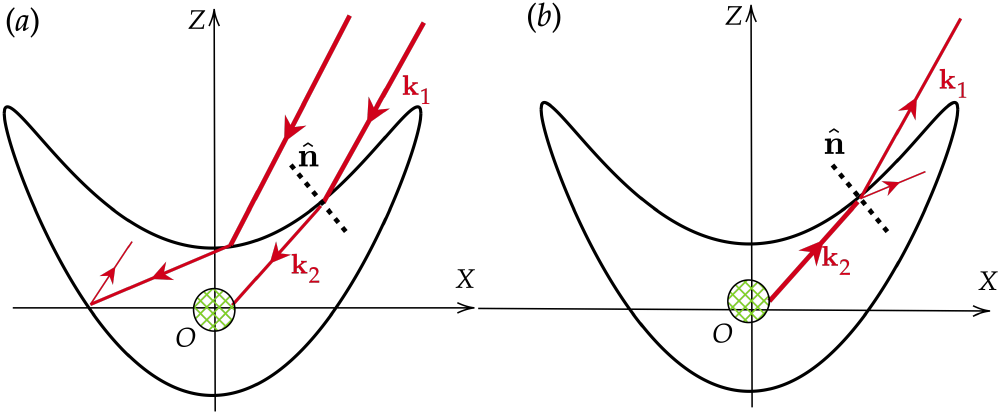
The two optical problems discussed here. Red lines are principal light rays. (a) The incoming problem of how cell shape influences intensity of photosynthetic light falling on a chloroplast. (b) The outgoing problem explores how the bioluminescence emission is influenced by the cell geometry.

### The outgoing problem: How cell geometry shapes bioluminescence emission

In marine environments, suspended dinoflagellates respond to the fluid flows associated with ambient turbulence and disturbances from large predators by giving off bioluminescent flashes. Figure 3(b) illustrates the mathematical problem of interest: how a cell’s shape alters the angular distribution of bioluminescent light emitted isotropically from small sources within the cell, undergoing both refraction and internal reflection. Since the cell is much smaller than the distance to predators, the focus is on the direction of emitted light rather than its origin.

**TABLE I.**
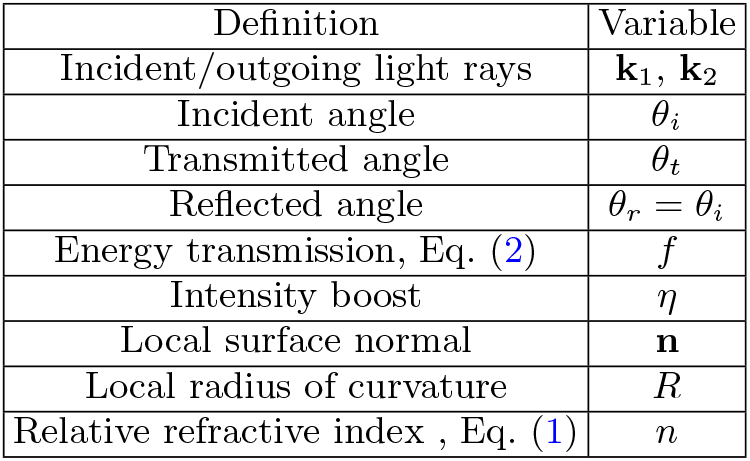
List of variables and their definitions.

| Definition | Variable |
| --- | --- |
| Incident/outgoing light rays | $\mathbf{k}_1, \mathbf{k}_2$ |
| Incident angle | $\theta_i$ |
| Transmitted angle | $\theta_t$ |
| Reflected angle | $\theta_r = \theta_i$ |
| Energy transmission, Eq. (2) | $f$ |
| Intensity boost | $\eta$ |
| Local surface normal | $\mathbf{n}$ |
| Local radius of curvature | $R$ |
| Relative refractive index, Eq. (1) | $n$ |

Our analysis assumes that the interior of a photosynthetic cell is a homogeneous optical medium with refractive index *n*_cell_, while the surrounding aqueous environment has refractive index *n*_water_ ≃ 1.33. For example, *Chlamydomonas* cells have been reported to exhibit *n*_cell_ ≃ 1.47 in the visible spectrum [2], which implies a relative refractive index *n* = *n*_cell_*/n*_water_ ≃ 1.1. We adopt this value in our calculations as representative of green algal exposed to light in aqueous environments.

To model how sunlight enters a cell and contributes to photosynthesis, we apply Snell’s law to incident light rays arriving from the surrounding water

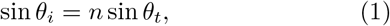

where *θ*_*i*_ is the angle of incidence in water, and *θ*_*t*_ is the angle of transmission inside the cell, both measured with respect to the local normal.

The fraction of energy transmitted into the cell quantifies how much unpolarized light passes through the interface. For incident angles *θ*_*i*_ *< θ*_*c*_ = sin^−1^(1*/n*), where total internal reflection does not occur, the Fresnel transmission coefficient *f* (*θ*_*i*_) for unpolarized light is [33]

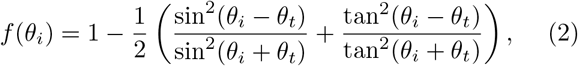

where *θ*_*t*_ = arcsin (sin *θ*_*i*_*/n*) is the transmitted angle determined by Snell’s law. Normal incidence (*θ*_*i*_ = 0) gives the maximum transmittance rate *f* (0) = 4*n/*(*n* + 1)^2^. The quantify 1 − *f* (*θ*_*i*_) is the proportion of light reflected back into the environment; it determines how much light penetrates the cell for internal processes.

We define the *boost factor η* as a measure of light amplification induced by the cell’s geometry. For the incoming problem, *η* = *d*Ω_2_*/d*Ω_1_ is the ratio of differential solid angles *d*Ω_*i*_ subtended at the entry and focus points, capturing how ray convergence enhances local light intensity. For the outgoing problem, *η* = *dA*_1_*/dA*_2_ expresses the relative change in projected beam area as light exits the cell, reflecting how geometry reshapes outgoing flux.

To simplify the analysis of light-ray interactions, we adopt the *chief-ray approximation*, in which all rays are assumed to deviate only slightly from a central principal ray. This reduces geometric complexity while preserving essential directional and focusing behavior.

#### Example: Optical Boost at the Algal Eyespot

We begin with a quantitative analysis of the results of Ueki, et al. [2] on the *eyeless* mutant of *C. reinhardtii*. Fig. 4 shows the geometry; a cell of radius *R* has a photo-sensor of radius *a* at its periphery, modeled as a circular patch. Light enters the cell from the opposite side, refracts at the surface, and the rays converge past the cell. A cone of these rays hits the photoreceptor. The maximum angular deviation of rays that reaches the photoreceptor defines a limiting incident angle *θ*_*i*_ with “impact parameter” *d* = *R* sin *θ*_*i*_ as in classical scattering theory.

**FIG. 4.**
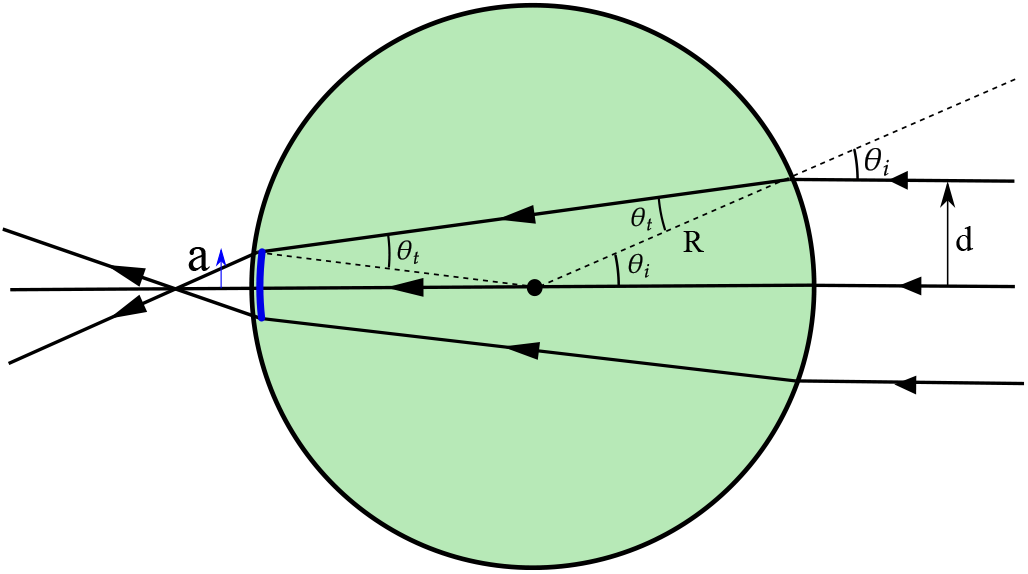
Analysis of the eyespot mutant in *Chlamydomonas*. The photosensor (blue) is located at the inner boundary of the cell. Although incoming light rays focus outside the cell, a gathering effect is experienced by the photosensor.

Figure 4 shows the geometry of interest. The vertex angle of the large isosceles triangle is *π* − 2*θ*_*t*_, so if we add all the vertex angles of triangles touching the center we find *a/R*+*π* −2*θ*_*t*_ +*θ*_*i*_ = *π*, and obtain a relation between the ratio *ϵ* ≡ *a/R* and the incident and refracted angles,

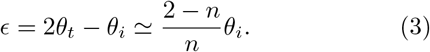

where the second relation follows from Snell’s law and is valid for *ϵ* ≪ 1, where a small-angle approximation holds.

The optical boost *η* is the ratio of areas of the incident light cone to that of the photoreceptor patch. In two dimensions, and in the small-angle approximation, this corresponds to the length ratio

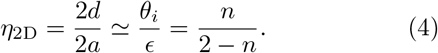

In three dimensions, the incoming rays span a circular disk of radius *d*, and the receiving region is a disk of radius *a*, so the boost becomes the area ratio

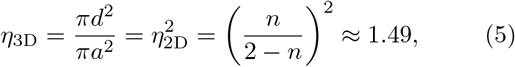

where we have used the value *n* = 1.1 for *Chlamydomonas*. This remarkably simple result shows that the index of refraction alone determines the boost for this simple spherical geometry. The quantity *η* − 1, the additional flux of light onto the photoreceptor, is

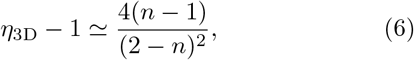

which is positive only when the relative index *n >* 1.

The large boost in Eq. (5) implies that an eyeless *Chlamydomonas* cell experiences two very distinct signals during each rotation about its body-fixed axis, the one from behind being 50% stronger than the other. Not surprisingly, that significantly larger signal dominates and the cell moves opposite to the wild type. The phototactic dynamics of eyeless mutants are discussed elsewhere [34].

## III. ANALYTICAL RESULTS FOR 2D BODIES

In this section we study the incoming problem for 2D bodies, progressing from simple to complex. We scale lengths by the size *R* of the body, so that the small target has radius *ϵ* = *a/R* and the bundle of light rays that intersects the target has half width *δ* = *d/R*.

### A. The Circle

We begin with the simplest case, a homogeneous circular body, which serves as a baseline for understanding how light refracts and concentrates within 2D bodies. Figure 5(a) shows the setup: a small, fully absorbing test ball of radius *ϵ* is at position (*r, φ*) where *r* ∈ [0, 1] is the (scaled) radial distance from the center and *φ* is the angle relative to the incoming light direction.

**FIG. 5.**
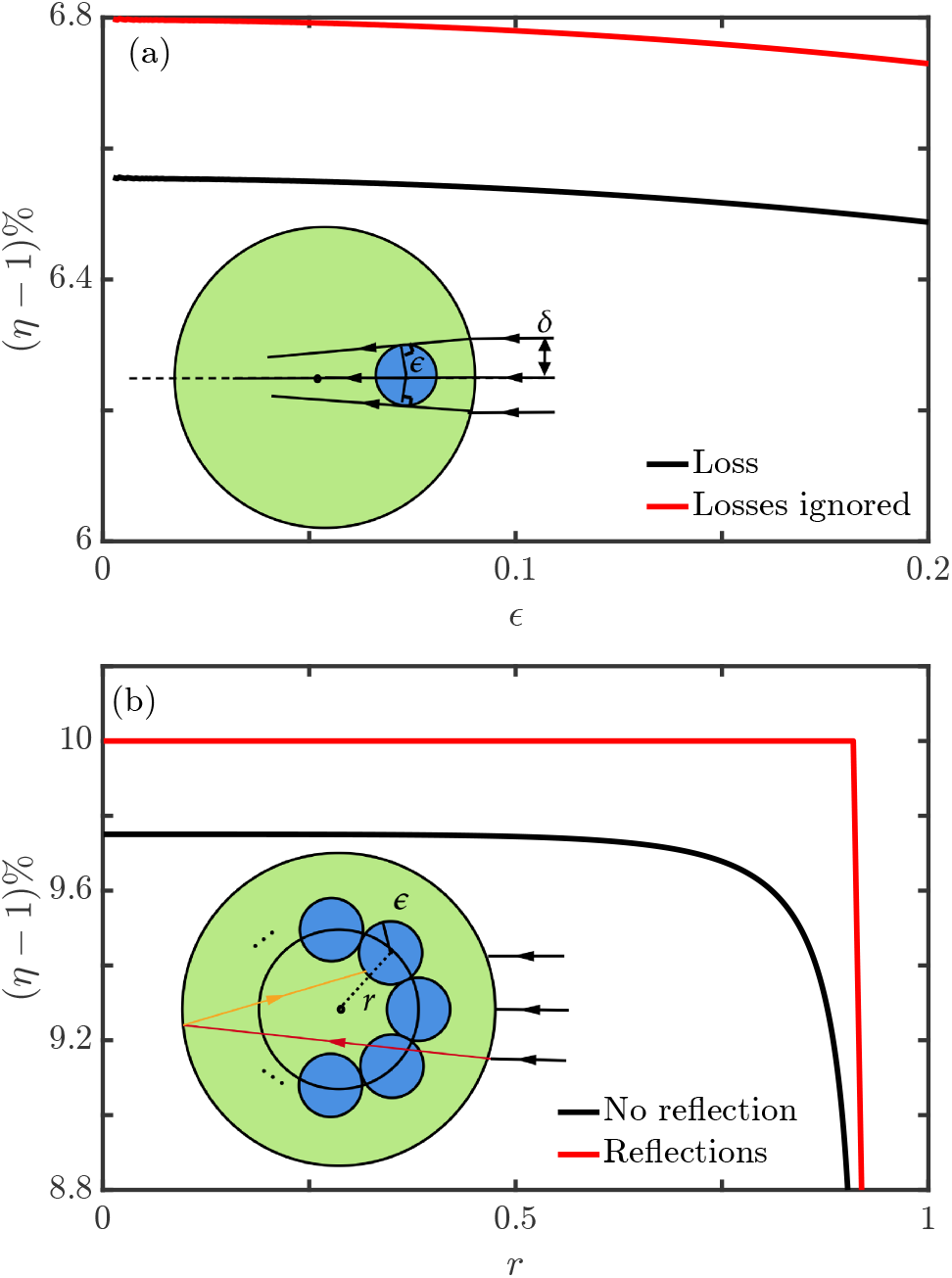
Incoming problem for a circle. (a) Boost versus ball size for *r* = 0.3 and relative index *n* = 1.1. As *ϵ* increases, *η* decreases, indicating weaker lensing and greater losses (2) when incident angle is large. (b) Disks of radius *ϵ* densely populate a circular shell. When multiple internal reflections are included, a ray (red) reaching the second disk is counted via the shell’s absorption. The boost is constant when all reflections (orange line) are included, until the ball reaches the total internal reflection zone where increasing *r* can no longer gain more energy. If reflections are excluded, *η* drops more near the upper boundary due to increased losses.

We first consider the case where the test ball is at the center of the circle to evaluate the maximal focusing effect. Because of the circular symmetry, we may take the incoming rays to be parallel and incident from the +*x* direction without loss of generality. The boundary entry points of rays that ultimately graze the edges of the test ball define the incident angular window. If a grazing ray enters at a point on the circle at angle *θ*_*i*_ then, since the normal to the circle at that point is radial, we have *δ* = sin *θ*_*i*_. And as the interior angle between the refracted ray grazing the target and the radial line from the origin that intersects the points of entry is *θ*_*t*_ we have the second relation *ϵ* = sin *θ*_*t*_, and thus *δ* = *ϵn*. Thus, the effective “window” of rays that can reach the test ball spans a lateral width *δ* that increases with the relative refractive index *n*. Note that the distance *δ* ≤ 1, so the corresponding boost *η*, the ratio of this width to the width of the test ball, is

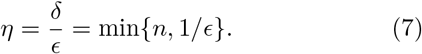

The boost equals the relative refractive index until it reduces because the ball can at most receive light with half-width *δ* = 1. The transition is also when total internal reflection occurs. This clean result captures the idealized case with perfect transmission and no optical loss, showing that the absorption at the center scales linearly with the refractive index *n*. In reality, partial reflection occurs at the boundary, especially for rays striking at oblique angles, due to refractive mismatch with the surrounding medium.

To incorporate such losses, we incorporate the angle-dependent transmission coefficient *f* (*θ*_*i*_) approximated using Eq. (2) for unpolarized light. The net boost factor, corrected for reflection losses, is then

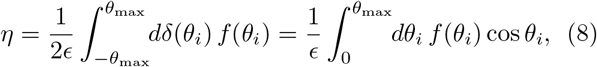

where *θ*_max_ = max {sin^−1^(*nϵ*), *π/*2} is the angular window of rays that refract to the center and cos *θ*_*i*_ is the Jacobian *∂δ*(*θ*_*i*_)*/∂θ*_*i*_. Eq. (8) reduces to (7) when *f* = 1.

### B. Effect of Absorber Size

We now consider test ball locations with *φ* = 0, so the geometry remains symmetric about the *x*-axis, but with *r >* 0. The boundary entry points of rays that graze the ball define a narrow angular range of incident rays. Using the geometry in Fig. 5(a), the condition for a ray to reach the disk is

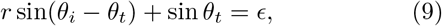

which implicitly defines *θ*_*i*_ as a function of *ϵ* and *r*. Once *θ*_*i*_(*ϵ*) is known, the boost factor *η* is calculated from Eq. (8). The resulting boost profile is plotted in Fig. 5(a) for *r* = 0.3. The boost is always smaller than the 10% boost when the ball is placed at the center, as in Eq. (7), because lensing is weaker closer to the light source. In the small-angle limit, (9) yields the boost

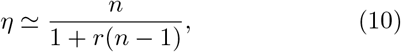

which interpolates between the limit *n/*(2 − *n*) in (4) for *r* = − 1, *n* for *r* = 0 and 1 as *r* → 1. The boost profile is continuous and converges to a constant value as *ϵ* → 0. This validates the use of small but finite test balls in simulations to estimate local light intensity. In biological terms, this corresponds to evaluating the light absorption by a small chloroplast placed at a given location within the cell. Because diffraction and coherence effects are negligible at this scale, the ray-based results are expected to agree with those from wave-optics models in the geometric optics limit.

### C. Angular Averaging

In natural biological contexts, light rays are likely to come from all direction relative to a cell. The results above should then be averaged of the distribution of incoming light rays, and the simplest assumption is a uniform distribution of the angle *φ* ∈ [0, 2*π*]. We can compute the boost at each angle, and perform this average, or equivalently, we can densely distribute *N* replicas of the test ball within a circular shell at radius *r*, with each ball interacting with light rays from different directions as illustrated in Fig. 5(b).

The circumference of the shell is the sum of the diameters of all the balls, so we have 2*ϵN* = 2*πr*. The total energy *E*_tot_ absorbed by the shell and the average energy *E*_avg_ absorbed by an individual test ball are related by

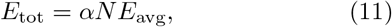

where *α* is inverse of the averaged number of times a light ray intersects a ball before reaching the cell wall.

We will show that the factor *α* = 1*/π* in two ways. First, note that the average energy absorbed is proportional to the circumference, as proved below in Eq. (34), so *E*_*tot*_*/NE*_*avg*_ = 2*πr/*2*πϵN*, indicating the result. Second, if we take the limit *r* → 0 while maintaining *ϵ/r* fixed to be a small number we are implementing a similarity transformation in which all ratios remain the same, and Eq. (11) still holds, with the same *α*. In this limit, all small balls approach the center, at which the total energy is the linear function *E*(*r*) = 2*rnf* (0), so

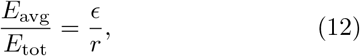

which then yields *α* = 1*/π*.

The light intensity profile as a function of the distance from the center of the circle is shown in Fig. 5(b), using the energy-averaging framework above. At each radial position *r*, the shell can be treated as a disk of radius *r* centered within the cell, and the local boost *η*(*r*) is computed using Eq. (8), with *θ*_max_ = max {sin^−1^(*nr*), *π/*2}. As *r* → 1, the shell approaches the boundary of the circle, and the transmitted light at large *θ*_*i*_ experiences greater loss when crossing the cell wall, as described by Eq. (2). This results in a decrease in *η*(*r*), illustrated by the black curve in Fig. 5(b), which drops sharply beyond *r* ≈ 0.75.

We extend this energy absorption analysis to include an arbitrary number of internal reflections f or the test ball. As before, the shell around the central disk is densely populated with smaller disks. These do not absorb light rays directly, as the presence of one does not interfere with another disk intercepting the same ray. Thus, we simply track the number of times a light ray passes through the shell. The total absorbed energy consists of the initial contribution from the ray passing through the cell wall, along with all subsequent internal reflections (see illustration in Fig. 5 (b), w here the thin red ray represents the first reflection). By symmetry, each reflected ray follows the same path as the initial one, preserving its angle of incidence.

Denoting the incoming light ray energy at angle *θ*_*i*_ as *E*(*θ*_*i*_), the total energy absorbed by the shell is

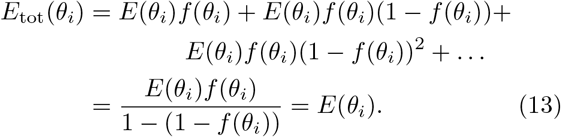

Thus, the contributions from all internal reflections exactly cancel the transmission losses, resulting in a constant boost given by Eq. (7), with {*η* = max *n*, 1*/r*}. This is illustrated by the red curve remaining flat until it drops significantly beyond *r* = 1*/*1.1 in Fig. 5(b).

### D. Duality for a Circle

In 2D, it can be shown explicitly that the incoming and outgoing problems are equivalent. We first prove this result for a circle without losses (2), and then proceed to the general case.

Consider first the outgoing problem. We adopt the notation ∠(**u, v**) for the angle between vectors **u** and **v**.

With reference to Fig. 6(a), the geometric boost for bio-luminescence *η*_*B*_ is defined as the limiting ratio of angles

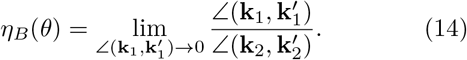

**FIG. 6.**
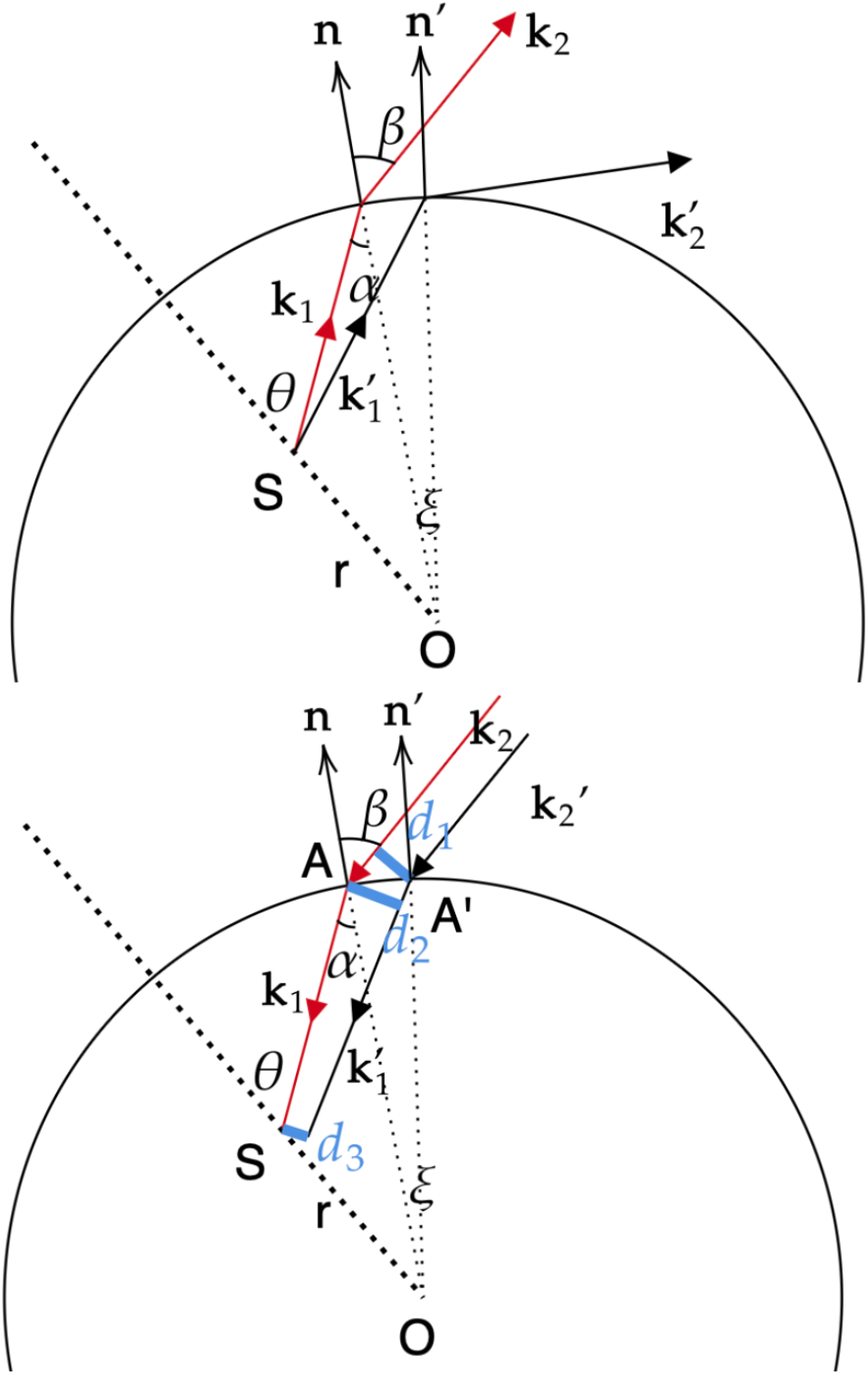
Geometry to demonstrate duality. (a) Bioluminescence case. Principal light rays are shown in red. We omit the angles between the light rays shown in black, denoted by *θ*^′^, *α*^′^ and *β*^′^, (b) Photosynthesis case. Incoming rays span a length *d*_1_, and the absorber has a projected length *d*_3_.

With *α* = ∠(**k**_1_, − **n**), *β* = ∠(**k**_2_, **n**), and *θ* = ∠(**k**_1_, **SO**), trigonometry and Snell’s Law (1) yield

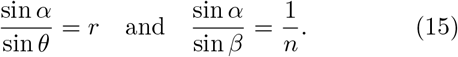

For the primed vectors and angles we have the analogous relations,

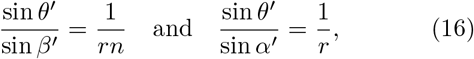

leading to relations for small changes in the angles,

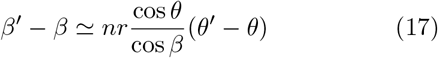

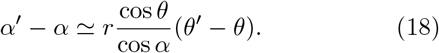

Let *γ* = ∠(**k**_2_, **OS**) = *θ* − *α* + *β* and similarly define *γ*^′^. Then from geometry we find

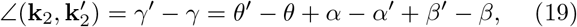

and hence, using Eq. (17) we obtain the boost (14)

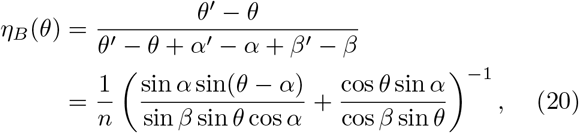

where in the last line we have used Eq. (15) and a product-to-sum trigonometric identity.

For the incoming (photosynthetic) case shown in Fig. 6(b), let point A denote the intersection of the refracted ray **k**_2_ with the cell wall, and similarly for A’.

The boost is the ratio of projected areas

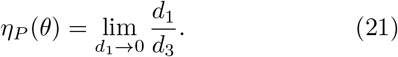

where *d*_1_ is the distance between the light rays **k**_2_ and 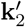, *d*_2_ is the distance between point A and 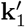. *S*^′^ is a point in the light ray 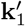 with **SS**^′^ ⊥ **SA**, and *d*_3_ = |**SS**^′^|. Also let 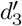 be the distance between S and 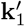.

Since *ξ* = ∠(**OA, OA**^′^) ≪ 1, we have ∠(**n, AA**^′^) ≃ *π/*2 and |**AA**^′^| ≃ *ξ*. Geometry then imposes the relations

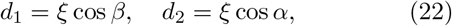

and

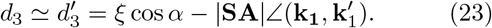

To work out the angle 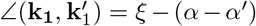, we apply Snell’s Law (1)

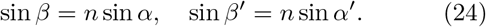

Similar to Eq. (17), we obtain

Combining with *ξ* = *β* − *β*^′^, we find

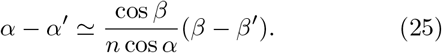

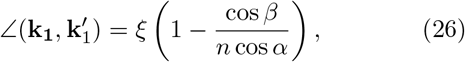

from which we obtain, using Eq. (22) and (23) in (21),

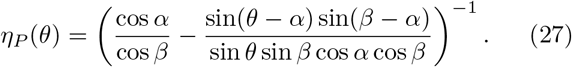

Trigonometry identities then lead to the duality result

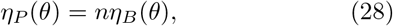

namely, the intensity profiles of the two cases are identical up to a factor of *n*. This is the two-dimensional analogue of “‘etendue” discussed below in § IV.

The outgoing light ray traces the full 2*π* angle:

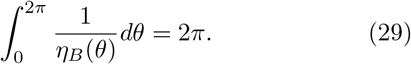

Then our definition (14) is *dθ/dγ* = *η*_*B*_(*θ*), leading to conservation of averaged photosynthesis boost,

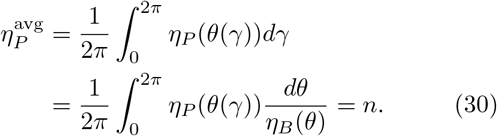

Here the light rays come at an angle *γ* with **OS**, uniformly distributed in [0, 2*π*].

### E. Duality for an Arbitrary 2D Body

For general shapes, any local region can be approximated as an arc segment with a specific radius of curvature. The argument applies universally for *r >* 1 (when the point lies outside the circle) or *r <* 0 (when the shape is concave). Consequently, the incoming and outgoing problems are locally and globally consistent, and (28) holds.

Note that for concave shapes, some radii of curvature may be negative, which implies that the local circle corresponding to that curvature lies outside the cell. In these cases, light rays tend to scatter rather than converge. Despite this, a similar analysis can be applied. If the test ball is placed at a distance *r* from the origin and has a small radius *ϵ*, the near-axis approximation holds, and the generalization of Eq. (27) is

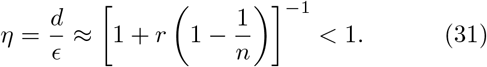

This result indicates that the test ball experiences a reduced boost, even before considering light losses.

In the presence of transmission loss, Eq. (29) represents energy conservation and applies generally, provided total internal reflection does not trap energy within the cell indefinitely. By the duality relation (28), Eq. (30) also holds universally: if no light path is confined solely by total reflection, the boost for the incoming problem remains unchanged. However, for structures like a circle, regions near the boundary experience total reflection, leading to a reduced average boost there. Likewise for the incoming problem, if the test ball is too close to the boundary, (7) leading to the reduced boost seen in Fig. 5(b). This issue is discussed further at the end of Sc. IV C.

### F. Averaging for 2D Shapes: A Surface Length Law

The baseline analysis of a circular test ball can be extended to arbitrary convex shapes, assuming isotropy in orientation—that is, the shape samples all orientations with equal likelihood. This condition is physically plausible for organisms such as algal cells undergoing slow tumbling or Brownian motion. Many unicellular organisms such as *Chlamydomonas* and *Volvox* [35] exhibit such dynamics due to flagellar motion or ambient fluid fluctuations. These lead to statistical averaging over orientations on timescales relevant for light exposure.

We parameterize such a shape as *r*(*θ*), with unit outward normal **n**(*θ*), and consider illumination from a fixed direction **k**_2_. The projected area is given by:

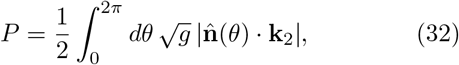

where *g* = *r*^2^(*θ*) + (*dr/dθ*)^2^ is the metric factor, and the prefactor of 1*/*2 reflects the fact that only the illuminated half contributes to the projection.

To compute the average projected area over all orientations, we rotate the shape by *θ*_0_ and average over *θ*_0_ ∈ [0, 2*π*]. Under such a rotation, the shape becomes

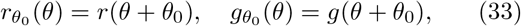

and the normal **n**_*θ*_0 (*θ*) is **n**(*θ*) rotated by *θ*_0_. Letting **n**^′^ = **n**(*θ*^′^), the averaged projection becomes

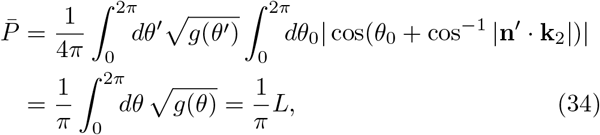

where *L* is the perimeter. Notably, the averaged projected area scales linearly with perimeter and is independent of shape geometry. This reinforces that projection-based absorption for fluctuating 2D convex bodies resspects a perimeter-based law.

## IV. ANALYTICAL RESULTS FOR 3D BODIES

The simplest shape 3D shape to analyze is the sphere. For the reference ball at the center, any incident light ray can be analyzed the same way as in 2D. The boost without loss of light due to reflection is analogous to Eq. (7),

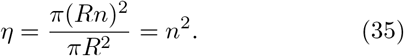

Considering loss, Eq. (8) becomes

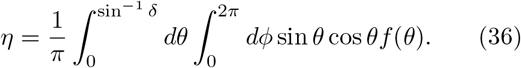

### B. Averaging in 3D: A Surface Area Law

The same principle discussed in § III.E extends to three-dimensional convex bodies with fluctuating orientations. Let the body have a fixed surface area and be parameterized by its radial function *r*(Ω), where Ω denotes a direction on the unit sphere, and **n**(Ω) is the outward unit normal. For a fixed direction **k**_2_ of incoming light, the projected area is

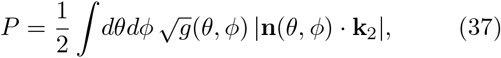

where the factor 1*/*2 counts the illuminated half of the body, and the metric factor is 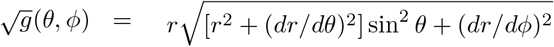

To compute the orientational average of *P*, we integral over all rotations Ω_0_. Without loss of generality, take **k**_2_ to align with the polar axis, and replicate the steps leading to (34), with *d*Ω_0_ = *dϕ*_0_*dθ*_0_ sin *θ*_0_, yielding

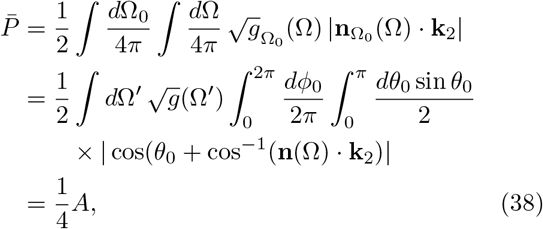

where *A* is the object’s surface area. As in the 2D case, this result depends only on total surface area. The average projected area of a convex 3D object under uniform orientation fluctuations thus follows a surface area law.

Also as in 2D, the above analysis breaks down for concave shapes. In those cases, parts of the surface can shade each other, leading to a reduction in the effective projected area. The impact of self-shadowing must be accounted for separately, as it can significantly alter the absorption characteristics and invalidate the simple surface-area scaling seen in convex geometries. We analyze such effects numerically in § V.

### C. Étendue and Flux Conservation

Étendue (often denoted ℰ) can be viewed as the “phase space volume” of a light beam [36]. It quantifies the spread of light in both position and direction. For a beam passing through a cross-sectional area *A* and contained within a solid angle Ω, the étendue is *E* = *A*Ω. In media where the refractive index varies spatially, this generalizes to

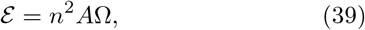

incorporating refractive effects on the direction of light rays. ℰ is conserved in passive optical systems, as we now show follows from flux considerations.

Assume a beam of light crosses the interface without loss and consider a corresponding set of rays defined in medium 1 by an area element *dA*_1_, with chief ray direction Ω_1_ and solid angle element *d*Ω_1_. After refraction into medium 2, these same rays will pass through some (generally different) area element *dA*_2_ and subtend a solid angle *d*Ω_2_ with direction Ω_2_. To connect *d*Ω_1_ and *d*Ω_2_, we must understand how a cone of rays in medium 1 maps into medium 2. From the differential of solid angle in spherical coordinates,

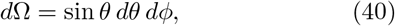

we differentiate Snell’s law and obtain

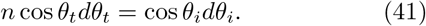

Since reflection does not change the azimuthal angle, we can multiply by *dϕ* on both sides and use Snell’s law again to obtain

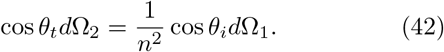

The light rays intersect the boundary in an area element *dA*, and if we define _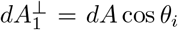_ and_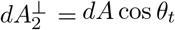_, we find

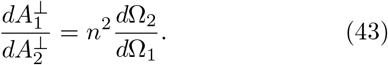

Under the chief ray approximation, each point in the area element has the same solid angle. The LHS of (43) is recognized as the boost factor for the incoming problem and the RHS as that of the outgoing problem. We thus establish that for each solid angle direction Ω_1_ for the outgoing problem,

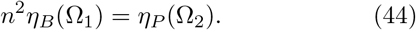

where *η*_*B*_ and *η*_*P*_ are boosts in the bioluminescence (outgoing) and photosynthesis (incoming) cases defined in (14) and (21). The duality in(44) is numerically verified in the next section, where the outgoing and incoming configurations yield equivalent intensity distributions.

In the same spirit as Eq. (30), we may calculate the average boost for the photosynthesis problem by integrating over incoming angles,

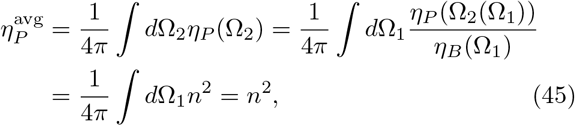

where in the second line we have used *η*_*B*_(Ω_1_) = *∂*Ω_2_*/∂*Ω_1_ as the Jacobian. The average boost is only a function of the relative reflective index *n*, and i s irrespective of the concave shell shape and the location of the test ball.

In fact, the argument above works for the case of multiple refractions, and when accounting for transmission losses as in Eq. (2). For the incoming problem, a principal light ray hits the test ball after the initial refraction and further reflections; we can trace back the ray from the test ball outside, which becomes the outgoing problem. The conservation of ‘etendue ensures that the duality of the boost hold at any point. The loss at the interface is also the same.

In considering all reflections and loss, let us label the boost by the number of reflections *i* that have occurred thus far. The outgoing boost is 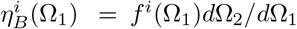, where 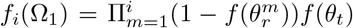 accounts for the loss according to Eq. (2). We similarly modify the definition of the incoming boost to account for multiple reflections, including losses. For the outgoing problem, all light will exit the body after a sufficiently large number of reflections, unless the situation is highly symmetric and some light rays are trapped by total internal reflection (See Fig. 9(b) where, near the edge, the average boost reduces to below 1). Then, energy conservation gives 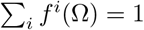, and thus

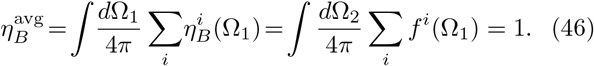

Since losses for incoming and outgoing light rays are the same in each direction, the (44) now reads 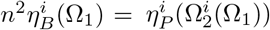. Denote the direction of the outgoing light ray after *i* reflections as 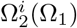. Eq. (45) becomes

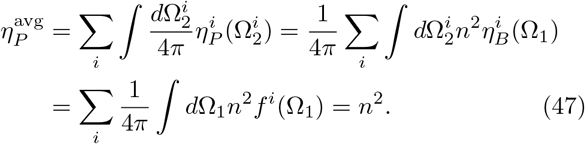

## V. NUMERICAL RESULTS FOR 3D BODIES

In this section, we present numerical results that support the conclusions drawn in the analytical sections above. We consider a two-parameter family of shapes for numerical computations. Built around the parameterization of ellipsoids, these shapes interpolate between the sphere (a suitable model for *Chlamydomonas*), and those with eccentricity approaching unity (appropriate to *P*.*fusiformis*) and then proceed to ellipsoids bent around their major axis (a shape like *P*.*lunula*). In a system of units made dimensionless by the semi-major axis *a* of the ellipsoidal limit, this family can be written as

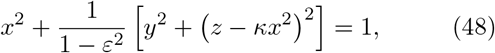

where 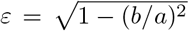 is the eccentricity and *b* is the semi-minor axis of the limiting ellipsoids obtained when *κ* = 0. These have major axes lying in the *xz*-plane with a radius *r*(*θ, ϕ*) measured from the ellipsoid center of

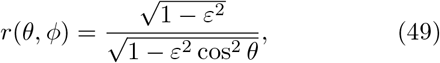

while for *κ* ≠ 0 the shapes are bent in the *xz*-plane around the curve *x* = *κx*^2^ for *x* ∈ [−1, 1],

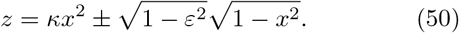

Figure 7 shows such shapes for various values of *ε* and *κ*.

**FIG. 7.**
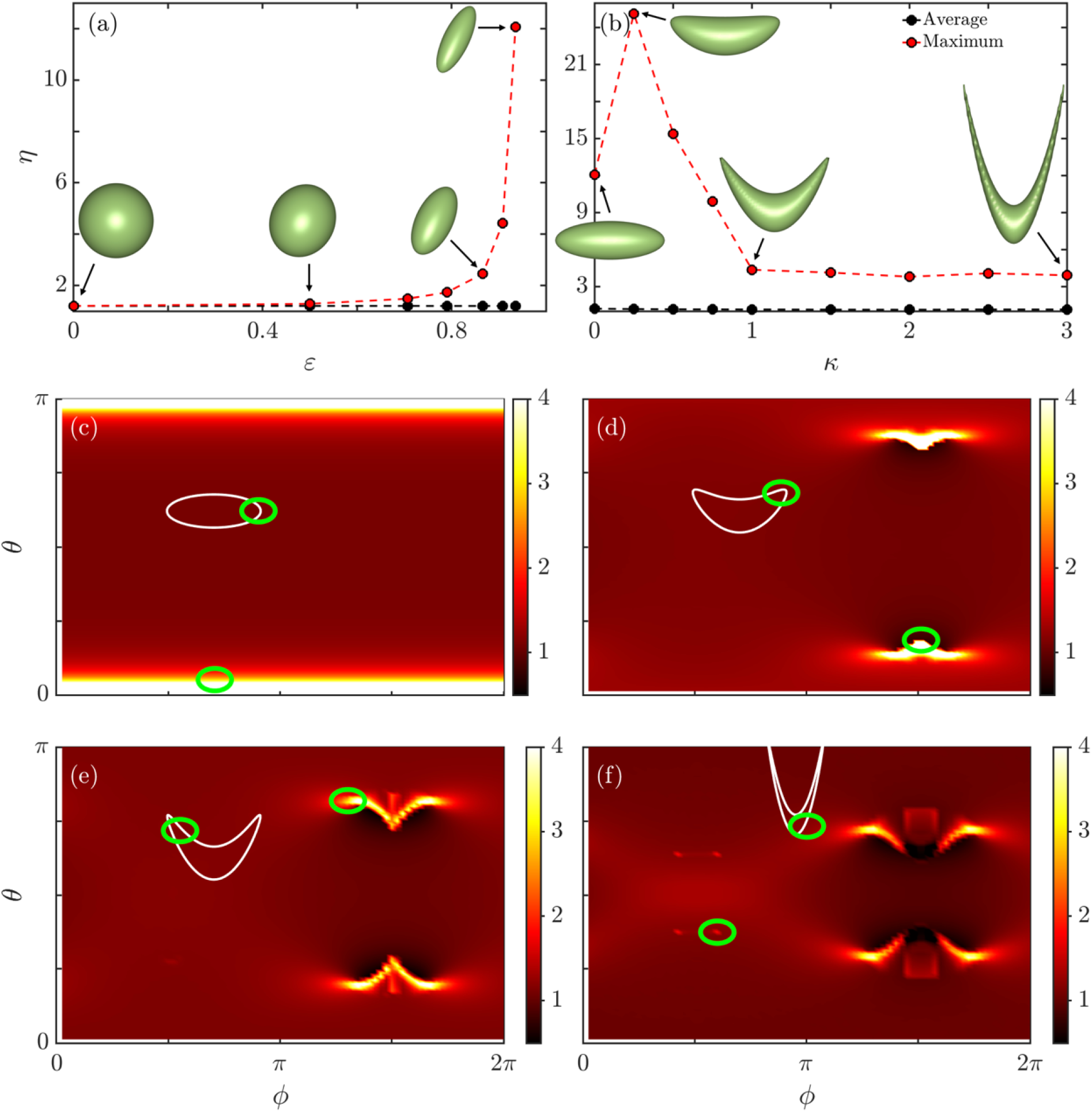
Variation with cell shape of maximum intensity boost at the center of the body. (a) Light intensity observed at the center of ellipses increases as a function of eccentricity *ε*, while the average intensity remains constant. (b) As a function of the curvature parameter *κ* the maximum boost for bent ellipsoids is nonmonotonic. (c-f) Intensity boost as a function of angles (*θ, ϕ*). For an ellipsoid, axisymmetry results in a boost that is independent of *ϕ*. Parameter values are *κ* = 0, 0.5, 1, 3 and eccentricity is 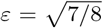. For bent shapes (d-f), the maximum stays near the direction of the tips. The green circles in (c-e) indicates the entry region giving strong boost. Note the focusing of the first reflections circled in green in (f).

### A. Photosynthesis at the Center

To examine how cell shape influences light concentration near the center, we model the chloroplast as a small, perfectly absorbing sphere at the center of the cell, shown as the green circle in Fig. 3(a). Incoming parallel light rays are assumed to arrive uniformly from all directions. We discretize angular directions and emit rays from a plane perpendicular to each specified propagation direction **k**_1_. Here and in following sections geometric symmetries allow us to restrict the angular sampling to half of the spherical polar angle domains in *ϕ* and *θ*. When the cell is bent, only two planar symmetries remain, resulting in boost-angle profiles with mirror symmetry across *ϕ* = *π/*2 (and 3*π/*2) and *θ* = *π/*2. Both *ϕ* and *θ* are discretized into 40 sampling points. For each direction, 360, 000 rays are generated so that the spacing between rays is smaller than the radius of the test ball, while ensuring that the ball remains small relative to the cell geometry.

Computational constraints restrict the region of ray entry to a zone centered on the test ball with a width 15 times its radius—an approximation that remains accurate, as confirmed by the close match between computed intensities in Fig. 7(a,b) and the analytical prediction Each ray carries equal energy, undergoes refraction at the cell boundary according to Snell’s law (1), and is attenuated by its Fresnel transmission coefficient (2). Each ray undergoes up to 11 reflections and those whose energy drops below 10% of their initial value are discarded to reduce computation time. This setup leads to billions of ray computations for each test ball position. The total absorbed energy is computed and compared to a case without a surrounding cell boundary.

We focus on two metrics: the maximum intensity boost (along the optimal direction) and the average boost (over all directions). In the spherical case, symmetry ensures that *η* is direction-independent, so the average and maximum coincide. As the eccentricity *ε* increases, Fig. 7(a) shows that a preferred direction emerges, with the maximum intensity increasing dramatically, from ∼ 1.2 to ~ 12. Introducing bending with a finite *κ* (Fig. 7(b)) further increases the peak, which exceeds 25 before declining to ∼ 4. This non-monotonic trend arises from the interplay between the increasing distance between the tip and the center, and decreasing radius of curvature radius at the tip. While the initial bending aligns focal zones with the center (by slightly increasing the axis length as initially the center lies between the focus and the tip), enhancing light concentration, further bending misaligns them, decreasing the central boost.

The average intensity is less sensitive to shape. For the sphere and ellipsoids, it remains near *n*^2^ ≃ 1.2 as predicted by (47). Bending decreases the average slightly, especially near the concave parts of the shell, due to redirection of light rays away from the center as *κ* increases.

### B. Angular Profile of the Boost

To understand better the entry point of the maximum boost and the distribution of light intensity, the boost observed at the center of the body is analyzed as a function of the incoming angles (*θ, ϕ*). For ellipsoids, the boost profile in Fig. 7(c) exhibits the expected axisymmetry in *ϕ*.

For mild bending, Fig. 7(d) shows that the peak region is centered at the tip of the bent shape and is the brightest spot in the profile. As bending increases in panels (e) and (f), light from the tip no longer focuses directly on the center, but rays entering from nearby directions do. This shift in the angular location of intensity peaks is accompanied by a migration of the *θ*-center of the bright regions, as the tip curvature becomes sharper.

As the shape becomes more extreme (panel f), additional bright spots and faint striping patterns emerge due to internal reflections of rays within the curved shell. The bright spots correspond to rays that enter the cell, reflect internally at the tip, and are redirected toward the center. The faint stripes are remnants of these multiply reflected paths. These reflection-induced features become especially prominent when the tip region is sharply curved and capable of directing incident rays back inward.

Dark regions in the angular map correspond to incident directions that either miss the central test ball or undergo reflection away from it. These regions typically have a boost near 1, indicating that the light reaching the test ball is neither enhanced nor significantly diminished.

### C. Spatial Distribution of the Boost

To gain further insight into the light intensity profile—and to explore the potential biological advantages of non-axisymmetric shapes and organelle distributions—we examine the spatial variation of light intensity within the cell. Specifically, we analyze both the average and the maximum intensity boost at various locations. The results are presented in Fig. 8. All plots consider only a smaller region inside the cell, as edge effects n ear the boundary make calculations unreliable.

**FIG. 8.**
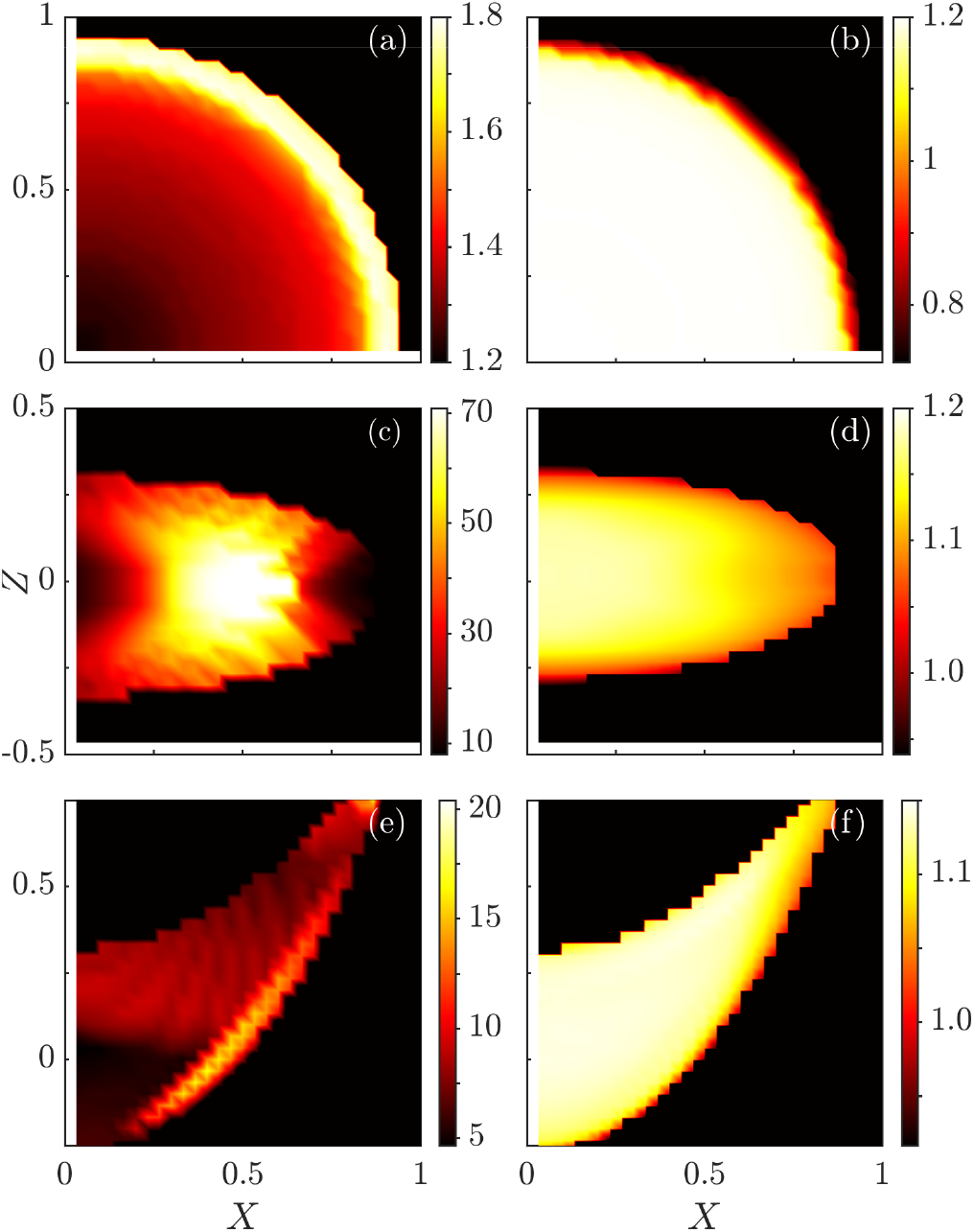
Spatial distribution of intensity boost. (a,b) The sphere ex bits a spherically symmetric profile. (c,d) Ellipsoid with 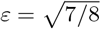. (e,f) *P. lunula* shape with *κ* = 1.

For the sphere, 40 sample points are taken along the radial direction, with the intensity distribution being invariant in the azimuthal angle *ϕ*. Therefore, discretizing the polar angle *θ* into 40 intervals is sufficient. For the ellipsoid, 208 sampling points are distributed inside the cell, leveraging symmetry along the *X*-axis. However, due to the reduced symmetry of this geometry, angular profiles a re only symmetric a bout *ϕ* = *π /*2. The high curvature of the ellipsoid surface demands a significantly finer angular grid; coarse sampling results in stripe-like artifacts—an issue commonly encountered in ray tracing simulations [41]. To eliminate these artifacts in the computed maximum intensity distribution, we discretize the angular space using 160 points in *θ* and 80 in *ϕ*. We use 8, 100 rays per direction, which has been found to yield accurate results in practice. For the *lunula* example, 387 sample points are used inside the shape, employing the same angular discretization and simulation parameters as with the ellipsoid. However, due to the fourth-order nature of the parametrizing equation (48), the simulation requires substantially more computational time.

Within the sphere, the averaged intensity is highly uniform, with a boost of ∼ 1.2 as discussed at the end of Sec. IV C. and as predicted by Eq. (47), until it drops to *<* 0.8 near the boundary. Variations are likely due to numerical errors and the fact that only finitely many reflected rays are considered, supporting the conjecture that all points within a convex shape experience the same average intensity. The maximum intensity is lowest at the center, increasing to ∼ 1.7 near the boundary. As the test ball approaches the boundary, the density profile becomes more uneven due to the proximity to the focal region of light from the antipodal point, as well as the increasing effectiveness of internal reflections. This is larger than the value calculated in Eq. (5) due to consideration of internal reflections.

A similar pattern is observed for the ellipsoidal case. Points near the focal region experience a maximum intensity boost exceeding 70, as light accumulates significantly near the focal point. The averaged intensity remains constant near the center, decreasing slightly from ∼ 1.18 (close to the expected value of *n*^2^) by ∼ 4%, then quickly drops to below 1, again likely due to the limited number of rays considered. For the *lunula* shape, the maximum intensity also shows sensitivity to discretization. However, the maximum boost is generally weaker than in the ellipsoidal case. The thin bright line in Fig. 9(e) results from a focusing effect at the tip. The average intensity remains nearly uniform near the center, decreasing from ∼ 1.14 to ∼ 1.1 near the tip due to the concavity of the shape. We also observe a decrease to below 1 near the boundary, similar to the ellipsoid case, likely due to limited light ray regions considered and the energy cutoff of light rays.

**FIG. 9.**
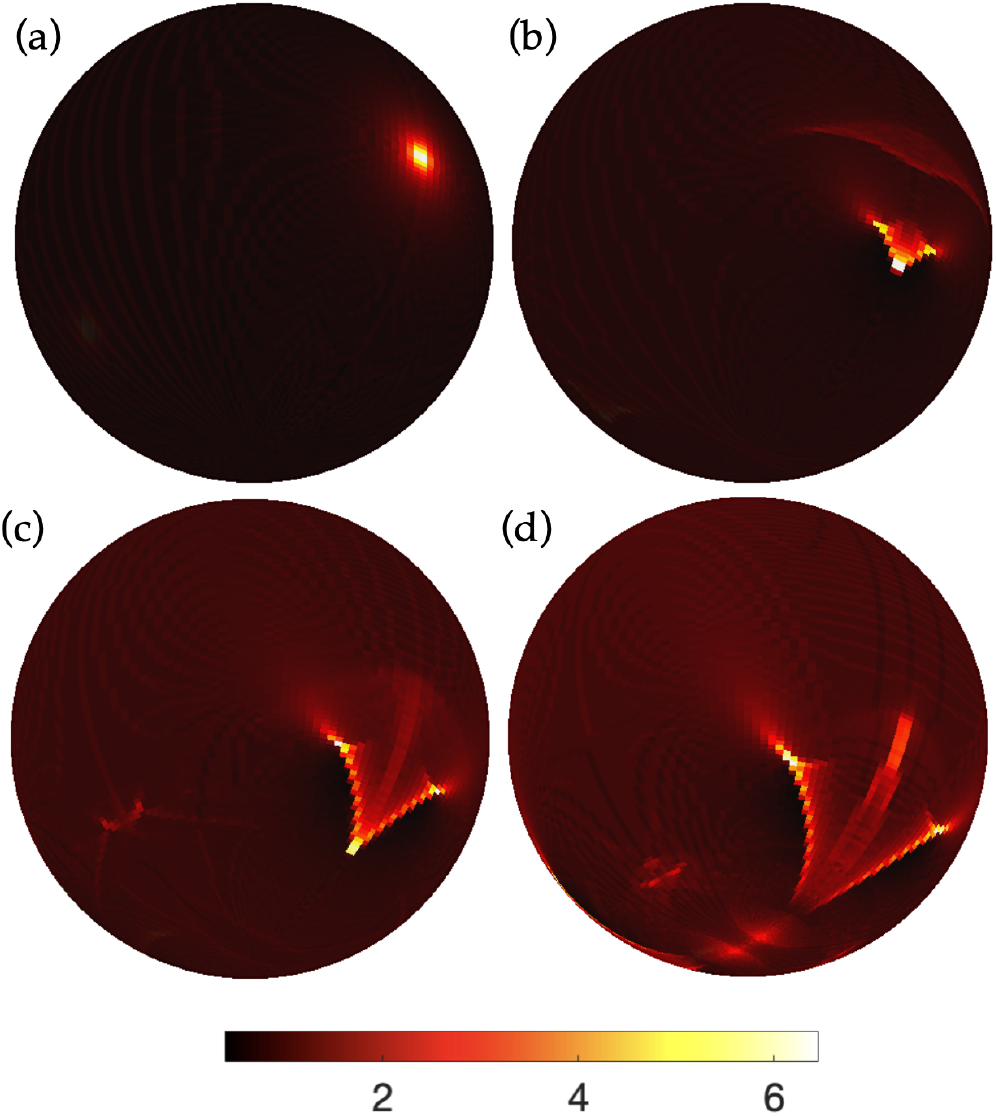
Angular distribution of light emission from the center of a l. For a source fixed at the center and with eccentricity 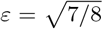 The bending parameter *κ* = 0, 0.5, 1 and 2 in (a-d). The boost region enlarges with *κ* while the maximum intensity first increases then decreases, a trend also seen in Fig. 7(b). The patterns resemble those of Figs. 7(c-f).

Overall, while the average intensity is robust across shapes, the spatial distribution of maxima reveals how cell geometry can create strong local variation. These focal hot spots highlight how even simple geometric differences can significantly modulate internal light fields.

### D. Bioluminescence

To model the outgoing problem, a point source emits light uniformly from a location inside a given cell shape. Light rays are refracted outward and reflected internally within the cell. Since the cell is small compared to the inter-organism distance, the location of outgoing rays can be ignored, and the light intensity distribution is evaluated on an infinite sphere centered at the cell, plotted with respect to the solid angle. The calculation is normalized against the case of a point light source at the center of the observation sphere. The computational process is simplified by generating light rays solely from the test sphere. The spherical grid is discretized into 160 × 160 points in *θ* and *ϕ*, respectively, and 409, 600 test light rays are generated from the ball. For reflections, we disregard those carrying less than 1% of their initial energy. The light intensity distribution at each location only requires millions of light ray realizations and achieves better angular resolution. Future studies on the spatial distribution of photosynthetic boost may benefit from employing bioluminescence-based simulation techniques. The light profile where the test body is placed at the center of various cell shapes is produced in Fig. 9, which shares the same pattern as Figs. 7(c-f). This further confirms the duality in Eq. (44).

We similarly examine the spatial distribution of the maximum intensity, as shown in Fig. 10. The maximum intensity values at each point are higher since, at the same angular resolution, the outgoing simulation converges more rapidly to the maximum. Notably, the bright regions remain consistent, in agreement with Eq. (44).

**FIG. 10.**
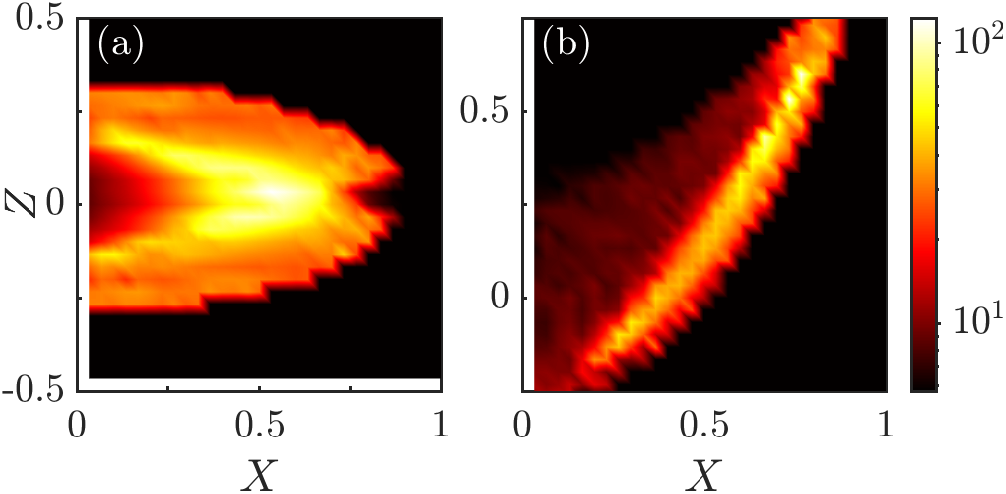
Spatial distribution of maximum focus boost. (a,b) correspond to (c,e) in Fig. 8. The plot is in log scale, with bright yellow regions showing strong focusing.

It is worth noting that our simulations of the incoming and outgoing problems handle concave shapes slightly differently. Specifically, light rays that re-enter the cell after reflection/refraction are not included in the analysis. These rays are expected to have negligible impact due to significant energy loss, as described by Eq. (2). Nevertheless, a more refined numerical treatment could be applied for concave geometries such as that of *P. lunula*.

## VI. DISCUSSION

We have presented a framework to understand how cell shape influences the distribution of light in organisms. Our work covers both incoming light, relevant for photosynthesis, and outgoing light, relevant for bioluminescence, and is based on geometrical optics. We introduced the notion of a boost factor *η* to quantify the focusing or defocusing of light within a cell. For simple geometries like the circle and the sphere, we obtained exact analytical results. In more complex shapes such as ellipses and bent geometries motivated by real dinoflagellates, we computed the intensity distributions numerically. In all cases, we find that geometric focusing alone can produce significant spatial variation in the light field, even in the absence of any specialized optical structures.

A key observation is that incoming and outgoing problems are related by a geometric duality, derived from conservation of ‘etendue in a passive optical system [36]. Duality allows us to deduce the intensity distribution in one case from that in the other. The duality relation was verified numerically across a range of geometries, and we showed that both configurations yield equivalent angular profiles after appropriate transformation.

Our results show that transparent, weakly refracting cells can experience substantial spatial variation in internal light intensity due to geometry alone. In bent cells, the light can be concentrated near the outer curvature by more than an order of magnitude. For convex geometries, the average boost is close to that of the circle or sphere, indicating that geometry tends to redistribute light rather than produce an amplification or reduction.

These results suggest that cells can passively manipulate light intensity using shape alone and may be relevant to how cells sense or respond to light. For photosynthetic cells, strategic localization of chloroplasts near focal zones (e.g., tips or curved regions) could maximize light absorption [37, 38]. For bioluminescent organisms, shaping the cell to direct emission could enhance signal directionality. For instance, our ray tracing simulations show that *P. fusiformis* emits light more strongly in the lateral directions than axially (Fig.2), perhaps confining in-plane signaling among neighboring organisms while reducing visibility from predators above or below, while *P. lunula* exhibits a concave-side bias and axial focusing, particularly when the source is centrally located. These anisotropies could influence signal projection, enhancing visibility in preferred directions while reducing the detection risk in others. Combining these directionalities with the inherent tumbling dynamics of elongated objects in shear flows, we may view cells as “stochastic beacons” of light, leading to nontrivial fluctuation statistics of light production in large groups of organisms. Recent studies of dinoflagellate light production and its cellular mechanisms suggest that geometric effects may also influence the triggering process leading to flashes [31]. Tools such as high-speed imaging or optogenetic reporters may provide ways to probe the internal light distribution in vivo [39]. Our findings may help explain why certain phytoplankton have evolved strongly eccentric or bent geometries, and they offer a testable hypothesis for sub-cellular organization driven by optical advantages.

Our model assumes geometrical optics and a homogeneous refractive index inside the cell. These assumptions are valid for cells much larger than the wavelength of light, but effects such as diffraction or interference may become relevant for smaller cells or near high-curvature regions. The presence of strongly absorbing components like pigments or organelles could also modify the results; the inclusion of absorption is an important future extension. Likewise, we have not modeled time-dependent effects such as cell rotation or deformation, which may alter the effective sampling of the light field. We also restricted attention to passive optical structures, neglecting any active light-guiding mechanisms or structural features. In some species, there is evidence of microlenses, reflective layers, or cellular-scale waveguides [40], which could be treated by extending the present framework to include multiple layers or graded refractive index profiles.

Finally, it is possible to couple the boost factor to light-activated processes, such as chloroplast migration or flash triggering. And the duality between incoming and outgoing problems may have implications for how phototactic and bioluminescent behaviors are coordinated in organisms that rely on both.

## ACKNOWLEDGMENTS

We are grateful to M.V. Berry for discussions at an early stage of this research, and to Mazi Jalaal for the image in Fig. 1(b). This work was supported in part by a Trinity College Summer Studentship and a Gates Scholarship (MY) and Grant No. 7523 from the Gordon and Betty Moore Foundation (SKB & REG).

The data that support the findings of this article are openly available [42].

## Notes

http://doi.org/10.5281/zenodo.14066435

### Competing Interest Statement

The authors have declared no competing interest.

